# A unified theory of E/I synaptic balance, quasicritical neuronal avalanches and asynchronous irregular spiking

**DOI:** 10.1101/2020.12.17.423201

**Authors:** Mauricio Girardi-Schappo, Emilio F. Galera, Tawan T. A. Carvalho, Ludmila Brochini, Nilton L. Kamiji, Antonio C. Roque, Osame Kinouchi

## Abstract

Neuronal avalanches and asynchronous irregular (AI) firing patterns have been thought to represent distinct frameworks to understand the brain spontaneous activity. The former is typically present in systems where there is a balance between the slow accumulation of tension and its fast dissipation, whereas the latter is accompanied by the balance between synaptic excitation and inhibition (E/I). Here, we develop a new theory of E/I balance that relies on two homeostatic adaptation mechanisms: the short-term depression of inhibition and the spike-dependent threshold increase. First, we turn off the adaptation and show that the so-called static system has a typical critical point commonly attributed to self-organized critical models. Then, we turn on the adaptation and show that the network evolves to a dynamic regime in which: (I) E/I synapses balance regardless of any parameter choice; (II) an AI firing pattern emerges; and (III) neuronal avalanches display power laws. This is the first time that these three phenomena appear simultaneously in the same network activity. Thus, we show that the once thought opposing frameworks may be unified into a single dynamics, provided that adaptation mechanisms are in place. In our model, the AI firing pattern is a direct consequence of the hovering close to the critical line where external inputs are compensated by threshold growth, creating synaptic balance for any E/I weight ratio.

**Highlights:**

- Asynchronous irregular (AI) firing happens together with power-law neuronal avalanches under self-organized synaptic balance.
- Self-organization towards the critical and balanced state (with AI and power-law avalanches) occur via short-term inhibition depression and firing threshold adaptation.
- The avalanche exponents match experimental findings.
- The adaptation time scales drive the self-organized dynamics towards different firing regimes.

**Author summary:** Two competing frameworks are employed to understand the brain spontaneous activity, both of which are backed by computational and experimental evidence: globally asynchronous and locally irregular (AI) activity arises in excitatory/inhibitory balanced networks subjected to external stimuli, whereas avalanche activity emerge in excitable systems on the critical point between active and inactive states. Here, we develop a new theory for E/I networks and show that there is a state where synaptic balance coexists with AI firing and power-law distributed neuronal avalanches. This regime is achieved through the introducing of short-term depression of inhibitory synapses and spike-dependent threshold adaptation. Thus, the system self-organizes towards the balance point, such that its AI activity arises from quasicritical fluctuations. The need for two independent adaptive mechanisms explains why different dynamical states are observed in the brain.

## Introduction

Self-organized quasicriticality (SOqC), as defined by Bonachela and Muñoz [1], appears in nonconservative models for forest-fires, earthquakes and neuronal networks [2]. The self-organization consists of a local load-dissipation homeostatic mechanism incorporated in a system that has an underlying critical point. In the context of neuroscience, this point is often a second-order absorbing state transition where neuronal avalanches present apparent power law scaling. These adaptive networks frequently include only excitatory neurons, and the homeostatic mechanism is implemented as short-term synaptic depression or, sometimes, as adaptation of the firing probabilities or thresholds of the neurons (see [3, 4] for reviews). Due to the empirical observation of neuronal avalanches [5–7], these models are considered to be good representations of the spontaneous activity of the healthy cortex [8].

Alternatively, when inhibitory neurons are included in the model, a globally asynchronous and locally irregular (also known as AI) regime of spontaneous firing emerges (see [9, 10] for reviews). Such activity is generated by a dynamic balance of inhibitory and excitatory inputs to the cells [11], where the net synaptic current tends to remain close to zero. The AI regime usually requires tuning of the parameters of the network, such as the thresholds or the excitatory-inhibitory (E/I) synaptic weight ratio [12]. Although neuronal avalanches were not found in synchronous-irregular (SI) states of E/I networks [13], we showed that synaptic balance may be achieved via the same homeostatic mechanisms that generate a quasicritical state [14]. In addition, E/I networks with weak excitation and inhibition tend to show neuronal avalanches away from the AI regime [15].

Here, we develop the theory of a self-organized E/I network, where inhibition homeostatically adapts according to short-term synaptic depression and thresholds adapt by increasing after each neuronal firing. We derive analytical expressions for the firing rates and compare them to simulations. Interestingly, our network robustly evolves in time towards a state close to the critical point, where: (I) synapses are balanced, (II) the firing pattern is AI; and (III) neuronal avalanches emerge and follow a power law. The regime where these three distinct phenomena coexist has never been reported, and they emerge, in our framework, due to the two independent mechanisms of homeostatic adaptation that leads to self-organized quasicritical fluctuations. Specifically, synaptic balance occurs for any choice of parameters, *i.e.,* regardless of any sort of parameter tuning. Neuronal avalanches that obey power laws appear when the homeostatic adaptation time scales are large enough (hundreds to a few thousands of milliseconds, hence biologically plausible).

We show that threshold adaptation plays a dominant role when compared to synaptic depression, even when both are subjected to the same time scales. Thus, the self-organized system tends to hover in an AI state where external inputs are dynamically compensated by the spike-dependent thresholds.

In a previous work [14], we showed that without the adaptation in synapses and thresholds, the E/I network studied here presents a typical transition into an absorbing state, causing the different firing patterns (AI, SI, and critical avalanches) to be separated in the phase diagram. We also showed that dynamic synaptic balance occur, but we did not prove it rigorously, and we did not test whether the AI state generated by SOqC also displayed power law avalanches.

Thus, all of these developments are done the current work: first, we introduce the E/I network and the simulation details in the model section. The Mean-field approximation presents the results previously obtained for the static model, and then introduces the novel results with the study of a mean-field map for the homeostatic system; we calculate the firing rates, characterize the critical point, and analytically determine the quasicritical self-organization conditions for the network (both for its firing rates and for its synaptic currents). Finally, we present the neuronal avalanches and scaling of the model and discuss its implications in the last two sections.

## The model

We study the excitatory/inhibitory network of [14], where each neuron is a stochastic leaky integrate-and-fire unit with discrete time step equal to 1 ms, connected in an all-to-all graph. A binary variable indicates if the neuron fired (*X*[*t*] = 1) or not (*X*[*t*] = 0). The membrane potential of each cell *i* in either the excitatory (*E*) or inhibitory (*I*) population is given by

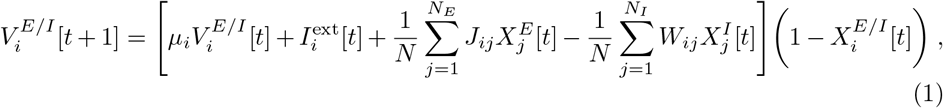

where *J_ij_* is the synaptic coupling strength, *W_ij_* = *gJ_ij_* is the inhibitory coupling strength, *g* is the inhibition to excitation (E/I) coupling strength ratio, *μ* is the leak time constant, and 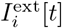 is an external current. The total number of neurons in the network is *N* = *N_E_* + *N_I_*, where the fractions of excitatory and inhibitory neurons are kept fixed at *p* = *N_E_/N* = 0.8 and *q* = *N_I_* / *N* = 0.2, respectively, as reported for cortical data [16] – although the values of *p* and *q* do not alter the dynamics of the network [14]. Note that the membrane potential is reset to zero at the *t* + 1 instant if a spike occurred at *t* because of the factor 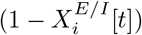.

All parameters are positively defined and are assumed to be self-averaging (i.e., single-peaked distributions with small coefficients of variation). Letting the excitatory synaptic current be 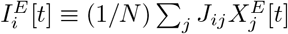 and the inhibitory be 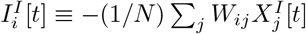, we define the net synaptic current,

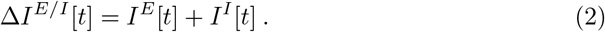

Eq. (1) is the same for both excitatory and inhibitory populations. This happens because the inhibitory self-coupling and the inhibitory coupling to the excitatory population are both given by *W_ij_*, whereas the self-coupling of the excitatory population and the coupling of the excitatory to the inhibitory population are both given by *J_ij_*.

At any time step, a neuron fires according to a piecewise linear sigmoidal probability Φ(*V*),

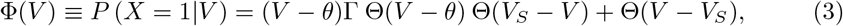

where *θ* is the firing threshold, Γ is the firing gain constant, *V_S_* = 1/Γ + *θ* is the saturation potential, and Θ(*x* > 0) = 1 (zero otherwise) is the step function.

The average over neurons of Eq. (1) yields the mean-field of this network. We call this the *static* version of the model, since the averages 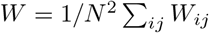 and *θ* = 1/ *N* ∑_*i*_ *θ_i_* are kept constant in time. It presents a mean-field directed percolation (MF-DP) critical point [14, 17] at *g* = *g_c_*, such that *g* < *g_c_* is the active excitation-dominated (supercritical) phase and *g* > *g_c_* corresponds to the inhibition-dominated absorbing state (subcritical). The synapses in the critical point *g_c_* are dynamically balanced: fluctuations in excitation are immediately followed by counter fluctuations in inhibition [14]. The avalanche distributions are the typical ones expected for a mean-field branching process. Moreover, the critical point can only be reached when the average external stimulus is of the order of the average threshold, *I*^ext^ ~ *θ*. Thus, we define the parameter *h* = *I*^ext^ − (1 − *μ*)*θ* as the *external suprathreshold current* ; it is analogous to an external field acting on standard models for directed percolation. The critical point is then (*g_c_, h_c_*) with *h_c_* = 0. A summary of the phase transitions undergone by the static model and its avalanches is given in Fig. 1.

**Fig 1.**
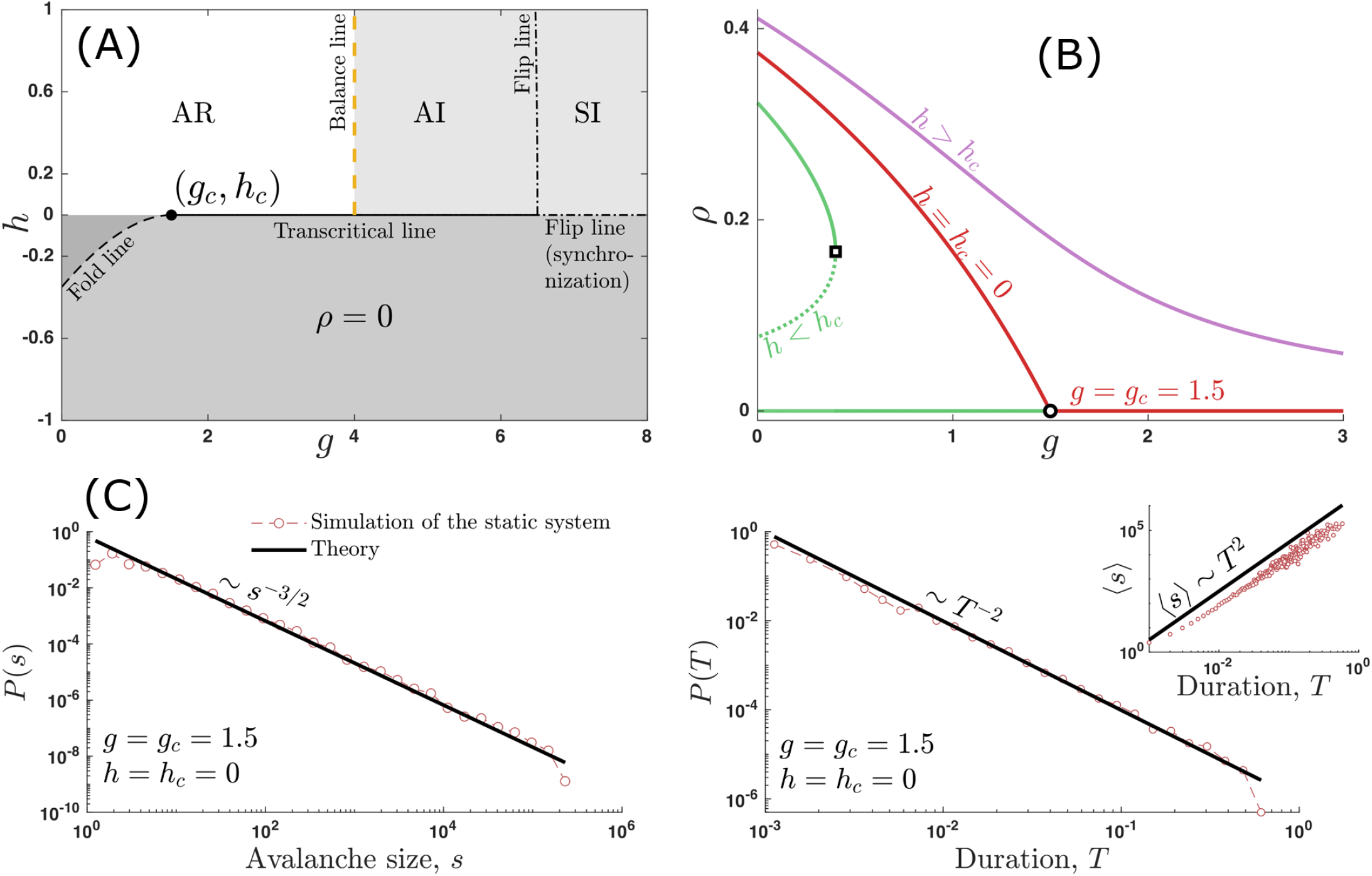
Phase diagram and avalanches of the static model. **(A)** and **(B)**: phase diagram given by the solutions of Eq. (19) with *μ* = 0, and *W* and *θ* constant in time. The axis parameters are given by: *g* = *W/J* and *h* = *I*^ext^ − (1 − *μ*)*θ*. The labels refer to the phases where the activity is: asynchronous regular (AR), asynchronous irregular (AI), and synchronous irregular (SI). The *ρ* = 0 is a phase with zero average firing rate. **(C)**: Avalanches on the critical point of the static model (constant *W* and *θ*) correspond to the expected MF-DP phase transition, yielding *P* (*s*) ~ *s*^−3/2^ and *P* (*T*) ~ *T*^−2^ [Eqs. (6) and (7)], with 〈*s*〉 ~ *T*^2^, respecting the crackling noise scaling relation [Eq. (8)]. The critical point (*g_c_, h_c_*) requires two fine-tunings, and it is separated from the AI state. Threshold adaptation will self-organize the system along the *h* axis and synaptic depression, along the *g* axis. In the text, we show that the addition of these homeostatic mechanisms are able to keep PL avalanches while generating a synaptically balanced AI state.

The hypothesis of brain criticality relies on the need for self-organization towards the critical point. However, the critical point requires two constraints, one on the E/I synaptic coupling *g* and the other on the balance of external currents by the threshold. We then introduce the *homeostatic* version of the model by letting *W* (the inhibitory coupling) and *θ* vary with time:

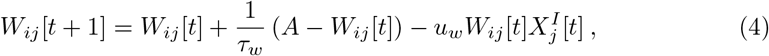

characterizing inhibitory depression, and threshold adaptation,

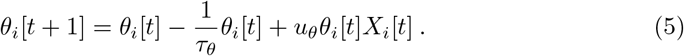

The parameters *u_θ_* and *u_W_* are the fractions of threshold increase and synaptic depression, respectively, *τ_θ_* and *τ_W_* should be moderately large decay time scales and *A* is a baseline synaptic coupling strength. Instead of inhibition, we can also homeostatically regulate the excitatory synapses [14], *J_ij_*, but experimental evidence suggests that it is the inhibitory synapses that tend to adapt to the excitation level of the network, and not the contrary [7, 9].

MF-DP systems are known to generate a SOqC state when excitatory synapses are depressing through an equation similar to the one above [2, 18]. Notwithstanding, here we have not only one, but two simultaneous homeostatic dynamics acting to self-organize the network. In a previous work, we concluded that this homeostatic version of the model is capable of generating a SOqC state possessing synaptic balance [14], but that conclusion relied only on numerical evidence. In the following sections, we will derive an analytical mean-field approximation for both the static and homeostatic versions of the model, and we will study the conditions for the emergence of the SOqC state in this network. Not only that, but we will also determine that the homeostatic model is always synaptically balanced, regardless of any parameter tuning. And finally, we will show that the SOqC dynamics generates power law (PL) avalanches when the activity is AI (as defined by a coefficient of variation, CV, of the network activity greater than one [19]).

## Methods

### Simulations

We ran simulations corresponding to Eqs. (1) to (5), and compared them to the analytical results developed in the forthcoming sections. Unless otherwise stated, we fixed the parameters *p* = 0.8 (fraction of excitatory neurons), *q* = 0.2 (fraction of inhibitory neurons), *I*^ext^ = 1 (external current), Γ = 0.2 (neuronal firing gain), *J* = 10 (excitatory weight), and *u_w_* = *u_θ_* = *u* = 0.1 (the fraction of inhibition depression and threshold adaptation, respectively). The parameter *μ* is the leakage time scale, and we show in the Mean-field approximation section that using *μ* = 0 does not change the phase transition, and hence this value is used for simulations. The remaining parameters, *N* (total number of neurons in the network), *τ_w_* (recovery time constant of the inhibitory synapses), *τ_θ_* (recovery time of the threshold) and *A* (maximum dynamical inhibition) were systematically explored. Some simulations have *τ* ≡ *τ_w_* = *τ_θ_*. Fig. 2 shows the average activity of the network simulated as a function of *τ* for different *N*, compared to the analytical description derived in the next section.

**Fig 2.**
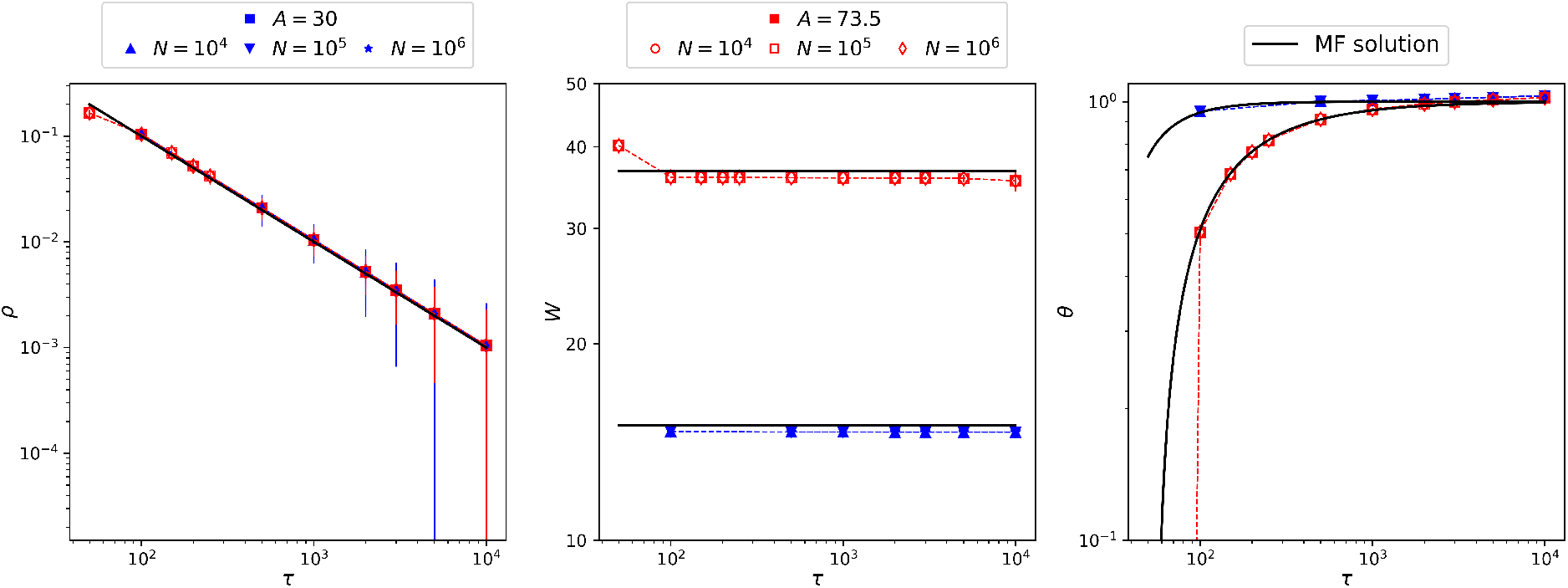
Network steady state activity. Average network activity as a function of *τ* compared to the mean-field analytic solution, Eq. (26) as a solid line. We ran simulations in two configurations: fine-tuned to the critical point, *A* = *A_c_* = 30 from Eq. (28) (▴ → *N* = 10^4^, ▾ → *N* = 10^5^, ★ → *N* = 10^6^), and in the supercritical regime, *A* = 73.5 > *A_c_* (○ → *N* = 10^4^, ◻ → *N* = 10^5^, ◊ → *N* = 10^6^). Symbols are the steady state averages of the network activity given by Eqs. (1)–(5); they fall on top of the MF approximation, except for *W* (*t*). The reason for this discrepancy in *W* is given in the text. Dashed lines between symbols serve only as guides to the eye.

All simulations were ran for a total of 1,000,000 time steps, and averages were calculated after discarding the first 10,000 time steps to avoid transient effects. We use the conversion factor of 1 time step = 1 millisecond, since a spike of a neuron, here, takes 1 time step.

### Avalanche computation and distributions

The simulation is the numerical solution of Eqs. (1) to (5), and we measure the instantaneous firing rate of the network with a total of *N* neurons, 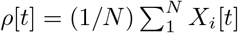, where *X_i_*[*t*] indicates a firing of neuron *i* at instant *t* (in any of the E/I populations). The total number of firings is *η*[*t*] = *Nρ*[*t*], and we choose the threshold *η*_th_ = *α*(max_*t*_ − *η*[*t*] min_*t*_ *η*[*t*]), with *α* = 0.2 as the fraction of the activity to be discarded. The results presented here are robust for a wide range of *α* values, provided that *α* is neither too close to zero nor to one. Then, we calculate the avalanches using the new time series, *η*′[*t*] = *η*[*t*] − *η*_th_: an avalanche of size *s_k_* and duration *T_k_* is defined as

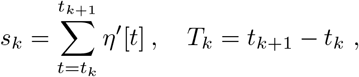

where *t_k_* and *t*_*k*+1_ are two consecutive moments of silence, i.e. *η*′[*t_k_*] = *η*′[*t*_*k*+1_] = 0 and 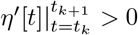. This method generates consistent scaling between 〈*s*〉 and *T* in the Ornstein-Uhlenbeck process [20], and also in the MF-DP avalanches (see Fig. 1).

We compare the distributions to the PLs

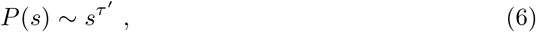

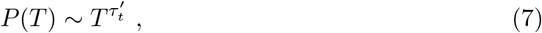

where we used a prime in the exponents to avoid confusion with the adaptation time scales, *τ*, *τ_θ_* and *τ_w_*, and keep consistent with the absorbing state phase transition literature [21]. When the system is critical, *s* and *T* are correlated through 〈*s*〉 ~ *T*^1/(*σν*⊥*z*)^, so the exponents *τ*′ and 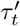 must follow the crackling noise scaling relation [21, 22]:

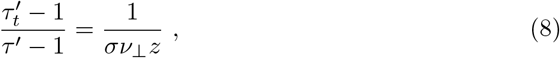

where *σ*, *ν*_⊥_ and *z* are other critical exponents. If the equality in Eq. (8) still holds after both sides being independently fitted, then it is a compelling evidence of a critical system. The left-hand side comes from the distribution of avalanches, and the right-hand side, from the 〈*s*〉 (*T*) plot.

### Mean-field approximation

The mean-field approximation is exact for our complete graph network with self-averaging parameters. Thus, we define the synaptic couplings as *J* = 〈*J_ij_*〉 and *W* = 〈*W_ij_*〉 (such that *g* = *W/J*), 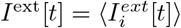 and *μ* = 〈*μ_i_*〉, where 〈·〉 means average over neurons. We also define the firing rates (the fraction of active sites per time step) 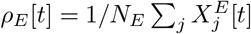 and 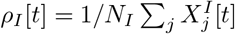. The fractions of excitatory and inhibitory neurons are *p* = *N_E_/N* and *q* = 1 − *p* = *N_I_ /N*, respectively.

Under these conditions, we can average Eq. (1):

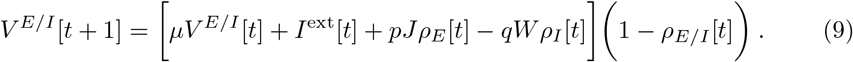

We may omit the *E/I* superscripts since this equation holds for both populations.

From the definition of the firing function, Φ(*V*) in Eq. (3), the model may have two types of stationary states: the active state, such that the density of active sites is *ρ_E_* = *ρ_I_* ≡ *ρ*\* > 0, and the quiescent state, where *ρ_E_* = *ρ_I_* ≡ *ρ*^0^ = 0. At instant *t* + 1, the active population is simply given by

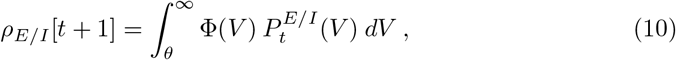

where Φ(*V*) is the conditional probability of firing at *t* + 1 given *V* at *t*, and 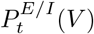 is the probability of having a neuron with potential *V* at time *t*.

This relation is sufficient for deriving the temporal evolution map for *ρ* because the reset of the potential causes the *k*-th subpopulation of neurons that fired together to also evolve together until they fire again. This allows us to write 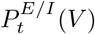 as a sum of subpopulations,

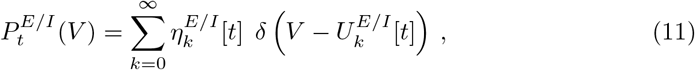

with *δ*(*V*) the Dirac’s delta function and

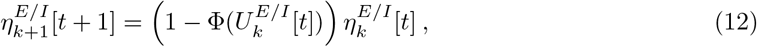

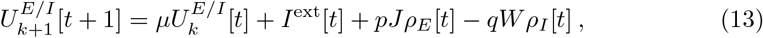

where 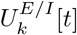 is the membrane potential of the *k*-th population of excitatory or inhibitory neurons that fired *k* time steps before *t* and 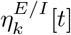 is the proportion of such neurons with respect to the total excitatory or inhibitory population. The potentials *U* come from Eq. (9), and the *k* = 0 population has 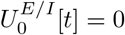 for any *t*, since the reset potential is zero. Then, Eq. (10) is reduced to:

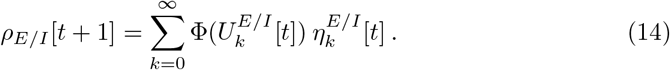

A neuron with firing age *k* at time *t* can, at time *t* + 1, either fire with probability 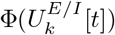 or become part of the population with firing age *k* + 1 that has density 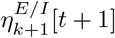.

The stationary state is given by the fixed point of Eq. (14). Inserting Eqs. (12) and (13) into Eq. (14) and assuming that the state of the system at *t* + 1 equals the state at *t*, we get:

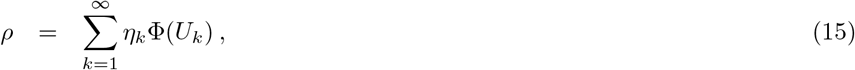

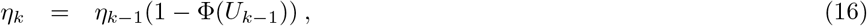

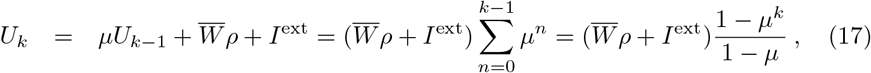

where *U*_0_ = 0, and we have used 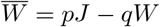 because *ρ_E_* = *ρ_I_* = *ρ*. Plugging Eqs. (16) and (17) into (15) and shifting the sum index yields:

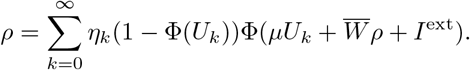

Considering the case where the stationary potentials lie on the linear part of the firing function, Φ, *i.e. θ* < *U*_1_ < … < *U*_∞_ < *V_S_*, with 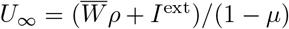 being the limiting potential when *k* → ∞. Then, using the expression from Eq. (3), we may explicitly write the second Φ(*V*) in the previous equation to get

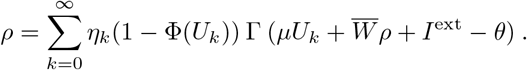

By using the relation in Eq. (15) and the normalization 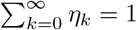, we obtain:

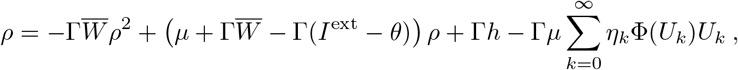

where we define *h* = *I*^ext^ − (1 − *μ*)*θ* as the suprathreshold external current, which may be regarded as an external field. Using equation (17) to describe *U_k_* only outside of the Φ function argument in the sum of the last term, leads to:

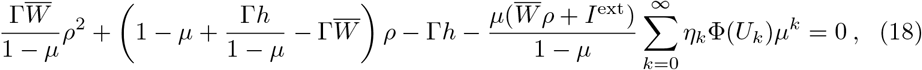

that holds when all stationary potentials lie in the linear region of the Φ function, i.e., when *U*_1_ > *θ* and *U*_∞_ < *V_S_*, as we have assumed (other solutions are trivial 2-cycles or a fixed point equal to zero – the absorbing state).

### Critical exponents without homeostasis

The leakage of the cells should not be instantaneous, but rather happen in the time scale given by ~ 1/*μ*. Hence, it is reasonable that 0 < *μ* ≤ 1 is the important range for the dynamics of the network. However, we show here that the value of *μ* does not change the critical exponents, and that *μ* = 0 is well-defined. Within this range, the sum in Eq. (18) converges to zero close to the critical point 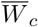, or whenever the network lies in a silent regime (zero firings in average). This happens because, when in silence, the firing rate of the network 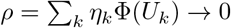 (which holds arbitrarily close to 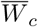 as well), and *η_k_*Φ(*U_k_*)*μ^k^* ≤ *η_k_*Φ(*U_k_*) for *k* ≥ 0. The firing rate *ρ* is the order parameter and, thus, it must go to zero on the critical point. In this case, Eq. (18) is simplified to:

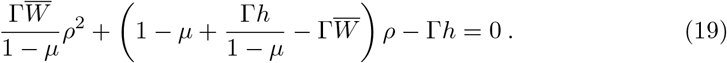

The critical point lies on the null external field, *h* = *h_c_* = 0, yielding the steady solution to Eq. (19), also known as the order parameter,

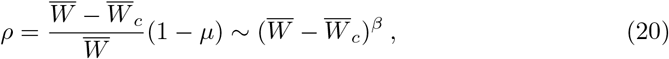

with *β* = 1 being its corresponding critical exponent, and

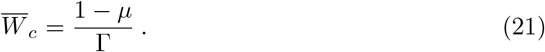

These relations can also be written in terms of the synaptic coupling ratio, *g* = *W/J*, as:

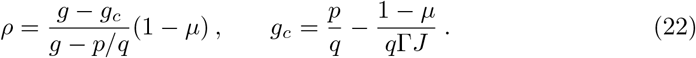

For *μ* = 0, the expression for *g_c_* reduces to the one obtained previously in Ref. [14].

The field exponent is obtained by isolating *h* in Eq. (19), taking 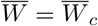, and expanding for small *ρ*,

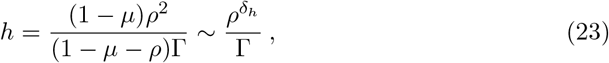

with *δ_h_* = 2. The susceptibility exponent, *χ* = *∂ρ/∂h* | _*h*=0_, is obtained by differentiating Eq. (19):

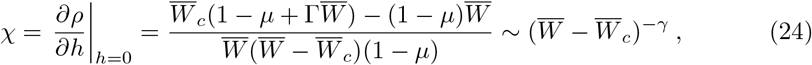

with *γ* = 1. This value, together with *β* = 1 and *δ_h_* = 2 puts this model into the mean-field directed percolation (MF-DP) universality class [21, 23] – see Fig. 1 for a complete phase diagram. Thus, having 0 ≤ *μ* < 1 does not change the nature of the phase transition undergone by the E/I network, as expected [14, 17, 24]. Also, we see that taking *μ* = 0 is well-defined, since Eqs. (18)–(24) remain valid. For that reason, we use *μ* = 0 in all our simulations.

### Is the homeostatic steady state SOqC?

So far, we have seen the static version of the model, where all the synapctic weights *J* and *W*, and the thresholds *θ*, remain constant over time. It has an MF-DP phase transition. Now, we will introduce homeostatic mechanisms to *W* and *θ* in the hopes of generating a SOqC state [1, 3]. We will derive a mean-field map for the dynamics of the network subjected to homeostatic synaptic inhibition and threshold adaptation. Many studies have shown that a single homeostatic dynamics leads to hovering around the critical point [25–29]. Here, we investigate in detail whether two independent homeostatic dynamics are able to do the same. The discussion in this section is systematically explored in terms of the phase diagrams of the E/I network in Fig. 3.

**Fig 3.**
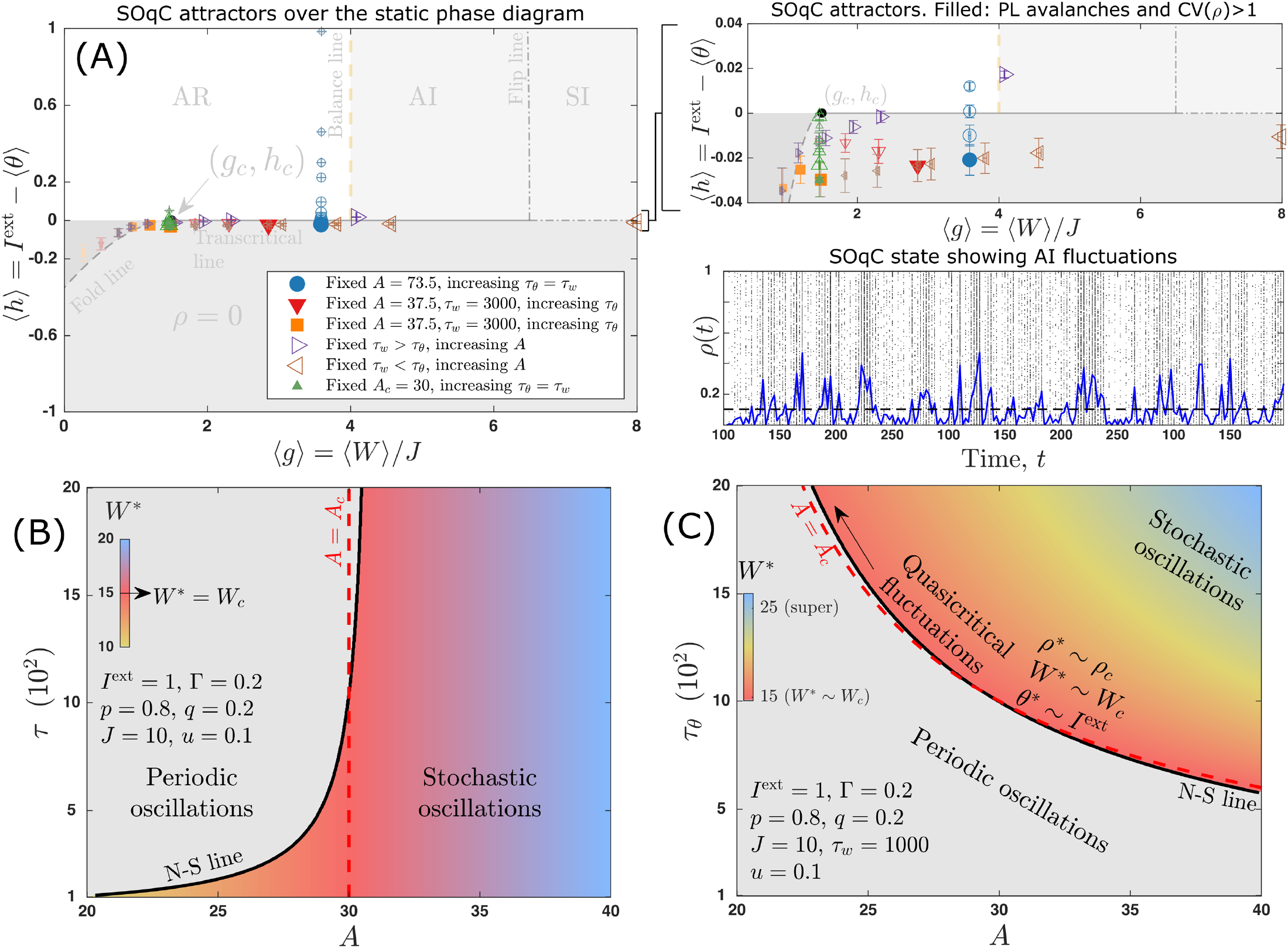
Attractors and phase diagrams of the homeostatic E/I network. **(A)** Simulations of the system given by Eqs. (1)–(5). The average of the steady state (i.e., the attractor) of each particular parameter configuration is plotted as a symbol. Increasing symbol sizes indicate the increasing of the parameter mentioned in the legend. Configurations that yielded AI activity with PL avalanche distributions are represented with filled symbols. AI activity is defined as having CV> 1 [19]. All parameter configurations yield dynamically balanced E/I synapses to 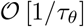 [Eq. (30)]. The attractors are overlayed on the static phase diagram to show that the network always converges to either: the bifurcation lines (Fold, transcritical and flip); or strictly to *h* ∼ *h_c_* = 0 when *τ_θ_* ≫ 1; but the synaptic depression is not able to lead the system towards *g* ∼ *g_c_* without tuning *A* = *A_c_*, even though PL avalanches are found in many different configurations with many *A*. Right panels: a zoomed-in version of the phase space in the left, and an example of a particular parameter configuration that yields AI and PL activity. Avalanches are calculated by thresholding the activity time series [20, 24]. **(C) and (D)** Phase diagrams of the homeostatic model in the plane *τ* vs. *A* (C) and *τ_θ_* vs. *A* for fixed *τ_w_* = 1000 (D). The solid line marks a numerically determined subcritical Neimark-Sacker (N-S) bifurcation from the stable fixed point of Eq. (25) (colored region), whereas the dashed line highlights the fine-tune, *A* = *A_c_* (configurations along this line correspond to the upward triangles in panel A). The color represents the value of *W*\*, so critical fluctuations, where the three variables, *ρ*[*t*], *W* [*t*] and *θ*[*t*], remain close to (*ρ_c_*, *W_c_*, *θ_c_*), are only found in the vicinity of the *A_c_* line (pink, red, orange colors). PL avalanches with dragon-king events (sporadic large avalanches) are found in the whole colored region for large enough *τ_θ_*.

Since *μ* does not change the critical exponents, we take *μ* = 0 for simplicity, such that all the subpopulations *k* ≥ 1 have the same membrane potential: from Eq. (13), *U*_*k*+1_[*t*] = *U_k_*[*t*] = *U*_1_[*t*] (recall that *U*_0_ = 0). This reduces the probability in Eq. (11) to *P_t_*(*V*) = *ρ*[*t*]*δ* (*V*) + (1 − *ρ*[*t*]) *δ*(*V − U*_1_[*t*]). Note that now there are only two subpopulations: the neurons that fired at time *t*, corresponding to *η*_0_[*t*] = *ρ*[*t*], and the neurons that did not fire, yielding *η*_1_[*t*] = 1 − *ρ*[*t*] from the normalization condition ∑_*k*_ *η_k_*[*t*] = 1.Using this probability in Eq. (10) and averaging Eqs. (4) and (5) over sites, we obtain the following 3-dimensional mean-field map:

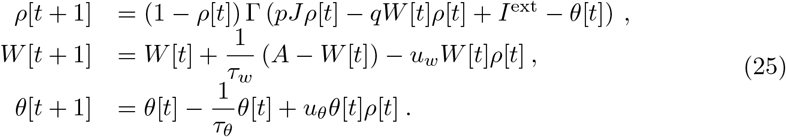

Here, we are only interested in the solutions with *ρ* > 0, since we used only the linear part of the Φ function to derive the *ρ*[*t*] map in Eq. (25). The steady state of the mean-field map is given by the fixed point

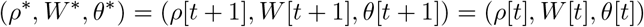

and yields two different types of solutions: the stable dynamics (when *θ* > 0),

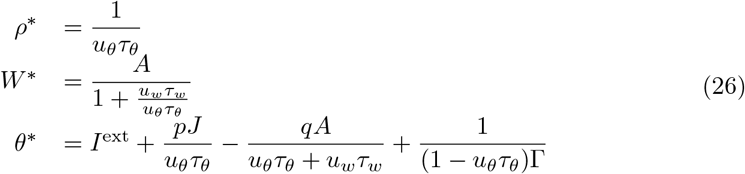

and the unstable hyper-excited state (*θ* = 0). The stability of the asymptotic solutions was numerically checked.

The critical point of the underlying system that we want to dynamically reach is given by 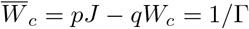 [Eq. (21)] and *h_c_* = *I*^ext^ − *θ_c_* = 0, which yields:

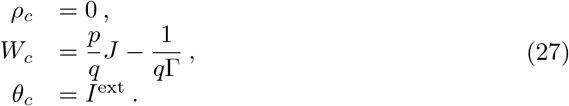

From Eq. (26), we see that taking fixed *u_θ_*, *u_w_* and *τ_w_* with *u_θ_τ_θ_* ≫ 1 is enough to achieve

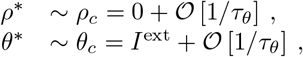

but that would still leave us with

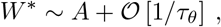

when *τ_θ_* ≠ *τ_w_*, or

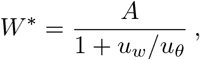

when both homeostatic dynamics have equal time constants.

Therefore, to meet the last requirement for reaching the critical point, *W*\* ≈ *W_c_*, we still need to fine-tune the parameter *A* = *A_c_*, with

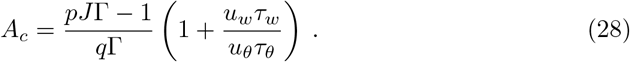

To understand why the fine-tuning is necessary, it is useful to isolate *W*\* from the *ρ*[*t*] map in Eq. (25), using *ρ*\* = 1/(*u_θ_τ_θ_*),

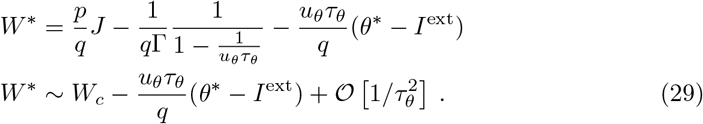

Since 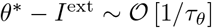, the last term in Eq. (29) is of 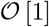, resulting in a displacement from the critical point along the *g* = *W*\*/*J* axis that is independent of *τ_θ_* (see Fig. 3).

Although the requirements upon *ρ*\* and *θ*\* are met through quasicritical fluctuations, the system still requires fine-tuning on the inhibitory synaptic amplitude. Such fine-tunings are known to be needed in self-organized neuronal systems without bulk-conservation of membrane potential [2]. They compensate for the dissipation of electrical activity by invariably forcing the synapses to restore themselves towards *A* after every spike.

However, an interesting result of having two homeostatic mechanisms is that Eq. (29) tells us how to overcome the fine-tuning: if 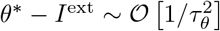, then all the requirements of homeostatic criticality would be met without any other parameter tuning. Conversely, Eq. (29) also means that the synaptic homeostasis is constantly being canceled by the threshold dynamics, avoiding the complete dynamical reach of the critical point. Nevertheless, the system is capable of self-organizing towards *ρ*\* ≈ 0 and *θ*\* ≈ *I*^ext^.

### Is the homeostatic steady state balanced?

The self-organizing of the system towards the critical point is weak, yielding a SOqC state due to dissipation. But what happens to the synaptic currents during this self-organization process? – here, we show that the SOqC state achieves perfectly balanced synaptic currents. This demonstration had only been done numerically in a previous work [14].

The net synaptic current is given by Eq. (2). In the mean-field steady state, we have the excitatory current *I^E^* = *pJρ*\*, and the inhibitory one, *I^I^* = −*qW*\**ρ*\*, where *ρ*\* and *W*\* are given by Eq. (26), resulting in the net contribution

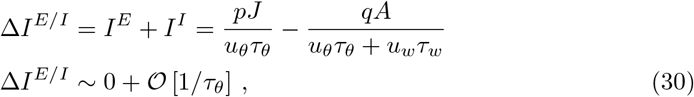

for fixed *u_θ_*, *u_w_* and *τ_w_*, with *u_θ_τ_θ_* ≫ 1. This analytical equation is in very good agreement with simulation results for the synaptic currents, *I^E^*, *I^I^* and Δ*I^E/I^* (Fig. 4). As *τ* and *N* grow, both the currents and the amplitudes tend to zero.

**Fig 4.**
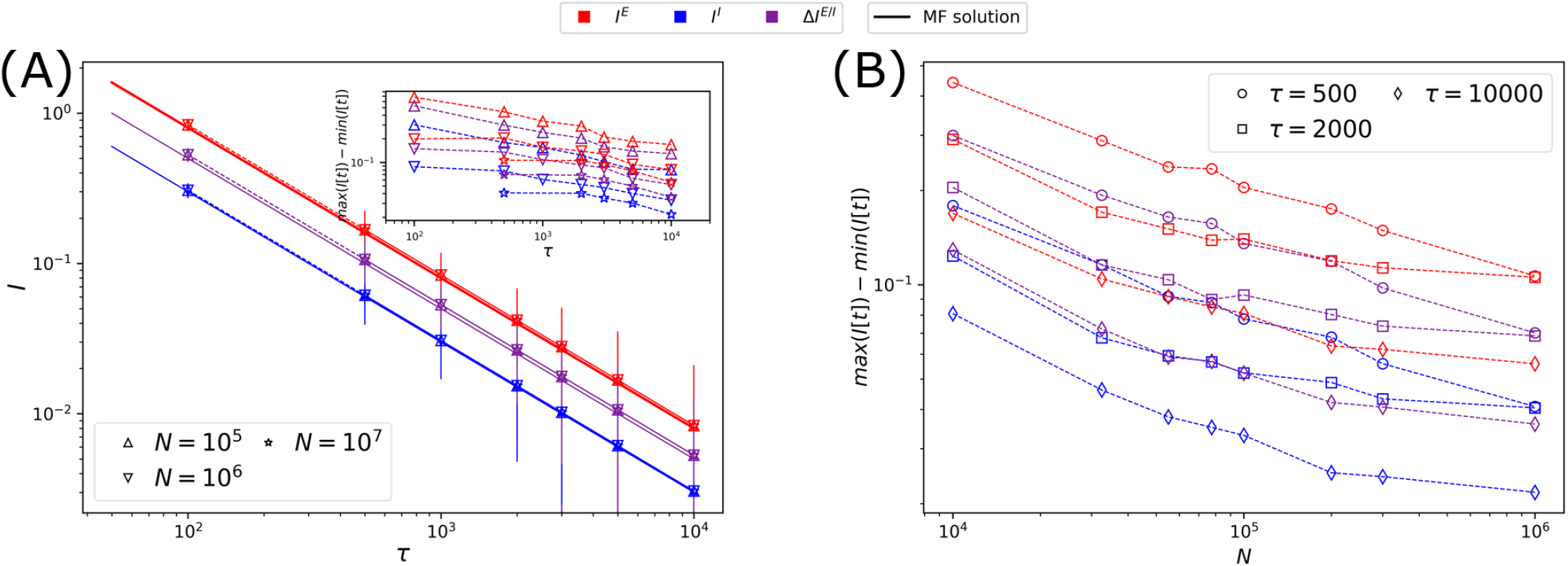
Synaptic currents. **(A)** Stationary value of the synaptic currents and their maximum amplitudes (inset) as a function of *τ*. Both tend to zero as *τ* grows, and the net steady current follows Eq. (30) with very good agreement. **(B)** The maximum amplitude of the synaptic currents decays as the total number of neurons in the network grows, meaning that the synaptic balance gets tighter with network size [10]. Symbols are simulation results and solid lines are the analytical calculations. Dashed lines between symbols serve only as guides to the eye.

This shows that the homeostatic system is balanced up to 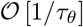. This means that although we may have different ratios of E/I currents (as expressed by *A/J*), and different regimes of steady activity, as evidenced by each attractor in Fig. 3, the system will still manage to dynamically balance its synaptic currents due to the homeostatic mechanisms.

### Avalanches and active cluster scaling

In Fig. 5, we show the avalanches of four particular configurations of the homeostatic dynamics: two with small *τ* and two with large *τ*. The addition of the homeostatic mechanisms creates a peak for very large (and very long) avalanches in *P* (*s*) and *P* (*T*), and breaks the crackling noise scaling relation (the fit of 〈*s*〉 vs. *T* yielded 1/(*σν*_⊥_*z*) between 1 and 2). For small adaptation time scales, the distributions are not PL – Fig 5(A). For large time scales (≳ 3000 ms) The exponents of the distributions are visually consistent with the MF-DP theory, but the maximum likelihood estimation [30] yields *τ*′ = 1.30 and 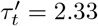 – Fig 5(B). The configuration with *A* = 73.5 and *τ* = 10^4^ in Fig. 5 corresponds to the filled circle in Fig. 3(A), hence it displays AI activity (since CV(*ρ*) > 1), PL avalanches and synaptic balance. The configuration with *A* = *A_c_*, however, displays PL avalanches without AI activity, since it is located in the periodic activity region of the phase diagram of Fig. 3(B).

**Fig 5.**
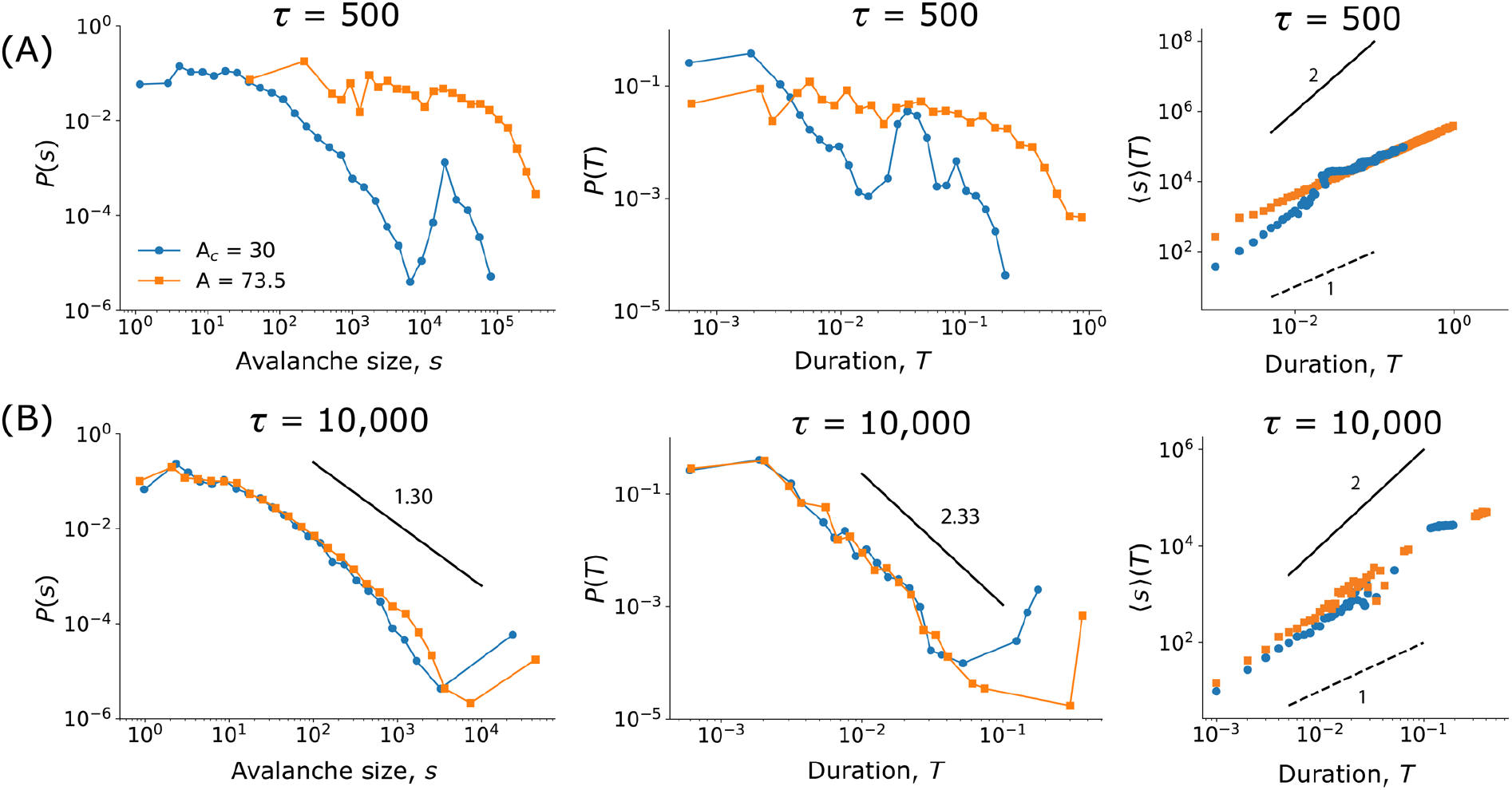
Avalanches of the homeostatic model. The distribution of avalanche sizes (*s*, left) and duration (*T*, center), together with the scaling relation (right), for *τ* = 500 (A) and *τ* = 10^4^ (B). Circles correspond to the homeostatic system fine-tuned to *A* = *A_c_* = 30 [Eq. (28)], whereas squares correspond to *A* = 73.5 > *A_c_*. The PL for *τ* = 10^5^ and *A* = 73.5 corresponds to the filled circle attractor in Fig. 3 yielding *P* (*s*) ~ *s*^−1.30^ and *P* (*T*) ~ *s*^−2.33^ (exponents obtained by maximum likelihood [30]). This particular configuration has all the features of E/I networks together: AI activity, synaptic balance and PL avalanches. The crackling noise scaling relation is not obeyed due to the SOqC dynamic. The solid and dashed lines in the 〈*s*〉 (*T*) plot (right column) serve only as guides to the eyes.

Beyond avalanche distributions, we also checked for the distribution *P* (*ρ*), since it is a direct measure of the network activity. Although not standard in the brain-related literature, some percolation models can be shown to obey a PL distribution for its instantaneous cluster sizes *P*_cluster_(*ρ*) ~ *ρ*^−*ϕ*^, where *ϕ* is known as the Fischer exponent [31]. This distribution is at times confused with the avalanche distribution, and has been shown experimentally to obey a PL in EEG and fMRI experiments [32, 33]. The static model has *ϕ* = 1 on the critical point *g* = *g_c_* with *h* = 0, and no PL in the super/subcritical regimes [see Fig. 6(A), top inset]. On the other hand, the homeostatic version of the model has *ϕ* = 2/3, and presents perfect collapse for different network sizes *N*. The same PL holds irrespective of *A* (Fig. 6) – a reminiscent evidence of the critical dynamics.

**Fig 6.**
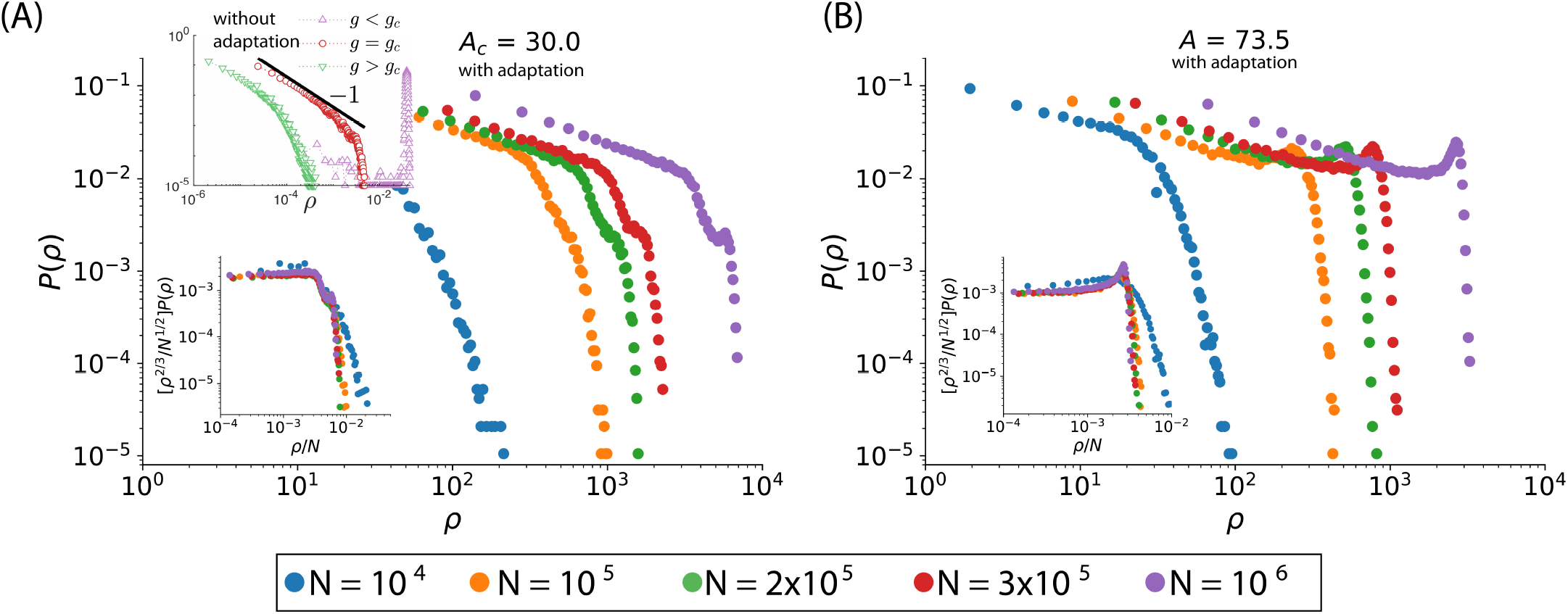
Distribution of active clusters. **(A)** Histogram of *ρ*[*t*] with self-organization (the homeostatic model, main panel) and without (the static model, top inset). For the static model, a PL is found *P*_cluster_(*ρ*) ~ *ρ*^−1^. Bottom inset: collapse of the data in the main panel, showing a scaling exponent *P*_cluster_(*ρ*) ~ *ρ*^−2/3^ for the homeostatic model at *A* = *A_c_*. **(B)** The distribution of active sites for *A* > *A_c_* yields the same exponent 2/3 as for *A* = *A_c_*, and also presents perfect finite-size scaling.

## Discussion

E/I networks are widely used as a theoretical model to understand neuronal activity: from memory dynamics in the healthy brain [34] to seizure development in epilepsy [35]. And absorbing state models can be employed to uncover features of the healthy [32] or unhealthy brain [36]. Amidst of that, there is an ongoing debate on the statistical features of the spontaneous activity of neuronal networks: should the network present asynchronous and irregular (AI) firing, or should it display neuronal avalanches in the form of activity bursts? Could both of these firing behaviors be reconciled? – In this paper, we offer a detailed theoretical study of a model that brings together both of these ideas: the quasicritical state (generated by the homeostatic mechanisms of synaptic depression and threshold adaptation) presents an AI firing pattern that is typical of E/I networks – see Fig. 3(A). Namely, the average activity fluctuates without a known periodicity, whereas the microscopic firing of neurons is irregular (any given neuron stochastically fires over time), such that the CV of *ρ*(*t*) is greater than one [19].

In addition, we showed that our homeostatic model displays PL-distributed avalanches that have size and duration exponents of about 1.3 and 2.33, respectively, determined by maximum likelihood. The only caveat is that these power laws do not agree with the PL coming from the 〈*s*〉 (*T*), as expected for a critical system. This is explained through the fact that the homeostatic dynamis can only generate a quasicritical state; and a quasicritical state is not able to display generic scale-invariance [1]. In our system as in typical SOqC neuronal networks [2], the critical point can only be dynamically reached with fine-tuning of the *A* parameter. The homeostatic dynamics is only enough to generate stochastic hovering around the critical line, but the system is never localized exactly at the critical point – the only spot in which the scaling must hold.

Interestingly, this is the first time we know of in which two independent homeostatic dynamics are shown to compete. Previous attempts of self-organization mainly focused on the adaptation of synapses (see [3, 4] for reviews). In this case, these systems also require tuning of parameters that are analogous to *A* in our model [2]. The extra adaptation mechanism presented here, given by threshold increments after each spike, makes our model more robust to changes in *A* when comparing avalanche distributions. However, the threshold time scale dominates the dynamics of our network and prevents a complete self-organization towards (*ρ_c_, W_c_, θ_c_*), since *W*\* − *W_c_* ~ *τ_θ_*(*θ*\* − *θ_c_*) and 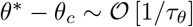. By letting *A* obey another adaptive mechanism, a phenomenon known as metaplasticity [37], we could potentially circumvent the need for tuning. Other solutions could involve having threshold adaptation that yield 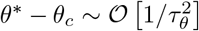 or altering the firing function, Φ(*V*), such that we get an equation for *ρ* that depends on *W* and *θ* through different types of nonlinearities.

The homeostatic adaptation of inhibitory synapses is inspired by experimental evidence [7, 9]. And here, together with threshold adaptation, it makes our network to always generate synaptic balance (to the 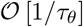, independently of any of the parameters of the model. This is another unifying aspect of our theory, since the balance happens concomitantly with the self-organized quasicritical fluctuations, where the model is visually consistent with complex activity patterns, expressed by the power law distributions of the activity (both for *ρ* and for *S* and *T*).

Looking for power laws in avalanche distributions is not the most reliable way of looking for a critical point. In absorbing state systems, such as branching processes, the theory gives us many other functions that are potentially observable (see [23], Table 4.1). In some models, the distribution of active clusters, *P* (*ρ*), is known to present a PL in the critical point [31]. Here, we showed that this is true both for the static and the homeostatic versions of the model. Although the cluster size exponent decreases from 1 to 2/3 with the addition of the adaptation mechanisms. PL cluster size distributions are also found experimentally in the brain [32, 33], although sometimes they are mistakenly referred to as avalanche distributions.

The quest put forward by the brain criticality hypothesis continues. Our model displays synaptic balance (independently of parameters) with AI firing patterns, under a two-variables self-organizing dynamic that drives the system close to the critical field, *h_c_* = 0; At the same time that this seemingly stochastic AI state takes place, the activity is complex: neuronal avalanches and activity clusters are distributed according to power laws – a reminiscent signature of the criticality that underlies adaptation mechanism. These features are typical of different fields in theoretical neuroscience, and our model is, then, a step towards putting these different frameworks together.

## Acknowledgments

This article was produced as part of the S. Paulo Research Foundation (FAPESP) Research, Innovation and Dissemination Center for Neuromathematics (CEPID NeuroMat, Grant No. 2013/07699-0). The authors also thank FAPESP support through Grants No. 2015/50122-0 (A.C.R.), 2016/03855-5 (N.L.K.), 2016/24676-1 (L.B.), 2018/09150-9 (M.G.-S.), 2018/20277-0 (A.C.R.), and 2019/12746-3 (O.K). A.C.R. thanks the financial support from the National Council of Scientific and Technological Development (CNPq), Grant No. 306251/2014-0. O.K. thanks the Center for Natural and Artificial Information Processing Systems (CNAIPS)-USP. M.G.-S. thanks the financial support of NSERC grant BCPIR/493076-2017 from A. Longtin and L. Maler.

